# Inhibition of histone lysine demethylase restores learning and memory in aged mice

**DOI:** 10.64898/2025.12.22.695858

**Authors:** Shail U. Bhatt, Lachlan MacBean, Enikö Kramár, Leticia Pérez-Sisqués, Phillip Smethurst, Josephine L. Robb, Neeru Jindal, Tracy L. Fetterly, Andrew Graham, Marcelo A. Wood, Sébastien Gillotin, K. Peter Giese, M. Albert Basson

## Abstract

Chromatin undergoes dramatic changes during the ageing process. In the brain, these chromatin changes are thought to underlie age-associated deficits in the activity-dependent gene transcription necessary for memory consolidation. Here, we show that the levels of a specific histone post-translational modification (PTM), trimethylation of lysine 4 on histone 3 (H3K4me3) is markedly increased in the hippocampus of aged mice and that the activity-induced increase in H3K4me3 that is observed in response to a learning stimulus in young mice, is severely blunted in the aged hippocampus. H3K4me3 typically marks open, accessible chromatin at the transcriptional start sites (TSSs) of actively transcribed genes. We identify altered H3K4me3 peaks at TSSs and show that ca. 90% of the activity-induced H3K4me3 changes at TSSs are either absent or reduced in the aged hippocampus. To understand the biological significance of these age-associated changes, we screened a library of pharmacological compounds for compounds that can alter H3K4me3 levels in hippocampal neurons. We show that treatment of aged mice with one of these, the LSD1 inhibitor ORY-1001, restored normal learning and memory in object location and recognition tasks. Furthermore, we show that ORY-1001 treatment increased long-term potentiation (LTP), a form of synaptic plasticity deficient in the aged hippocampus. These findings suggest that targeting the epigenetic machinery that regulates activity-dependent gene transcription may represent an avenue for treating age-associated cognitive impairment.

**Highlights:**

- H3K4me3 levels are elevated in the naive aged mouse hippocampus
- Activity-dependent H3K4me3 induction is impaired in the aged mouse hippocampus
- LSD1 inhibition restores learning and memory in aged mice
- LSD1 inhibition restores activity-dependent Arc induction and increases LTP in the aged hippocampus

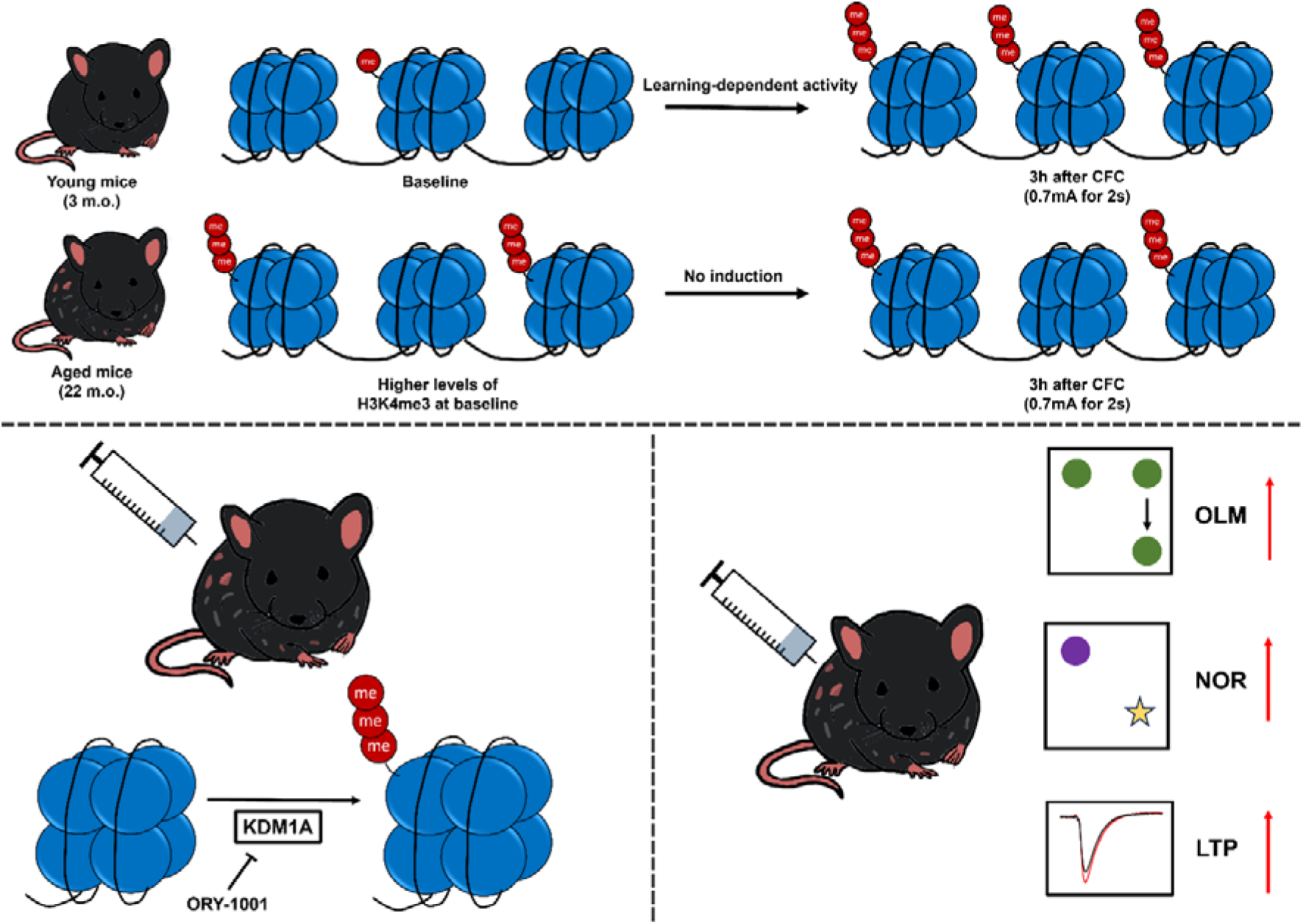

## Introduction

Mild cognitive impairment (MCI) represents a transition stage between healthy ageing and dementia, and a significant risk factor for Alzheimer’s disease, with >25% of individuals with MCI developing Alzheimer’s disease, compared to only 1% of those with normal cognitive functioning [1]. The hippocampus is particularly vulnerable to ageing, and age-associated changes in the hippocampus are thought to underlie deficits in associative learning and spatial memory [2]. Aged mice exhibit deficits in spatial [3–6], associative [7–8], and hippocampus-independent memory [9–10]. Studies on ageing rodents have identified potential mechanisms for age-associated learning and memory deficits, including neuroinflammation [11], abnormal microglial function [12] and epigenetic changes [13–14] all of which impair synaptic plasticity in the hippocampus.

Wide-spread epigenetic changes are characteristic of ageing cells [15]. Specific changes in histone acetylation have been reported in the aged hippocampus and targeting the histone acetylation machinery can reverse hippocampus-dependent learning and memory in aged animals [16–17]. For instance, deleting or inhibiting the histone deacetylase HDAC3 can restore learning and memory in aged mice [5, 18] and treatment of aged mice with the histone deacetylase (HDAC) inhibitor SAHA can restore activity-induced histone 4 lysine 12 acetylation (H4K12ac) in the aged hippocampus and improve hippocampus-dependent learning and memory [14].

Histone lysine methylation is a pivotal regulator of memory consolidation [19–20]. Studies in rats suggest that trimethylation of lysine 4 on histone 3 (H3K4me3), a modification found at active gene promoters, is induced in the hippocampus by a contextual fear stimulus [21], implicating activity-induced H3K4me3 in learning and memory. Collins et al. found that broad H3K4me3 peaks are associated with important learning and memory genes in the hippocampus [22] Mice heterozygous for *Kmt2a*, a gene encoding a histone lysine methyltransferase responsible for H3K4me3, exhibit learning and memory deficits. Furthermore, mice lacking either *Kmt2a* or *Kmt2b* [23] from excitatory neurons exhibited hippocampus-dependent learning and memory deficits [23–24]. H3K4me3 dysregulation has been linked to ageing in various organisms including yeast [25], C. elegans [26] and humans [27]. We therefore investigated the regulation and function of H3K4me3 during hippocampus-dependent learning and memory in mice and tested the hypothesis that changes in H3K4me3 underlie learning and memory deficits in aged mice. We established a high throughput screening platform to identify pharmacological compounds that can alter H3K4me3 levels in hippocampal neurons and evaluated the ability of one of these compounds to retore learning and memory in aged mice.

## Method details

### Experimental details

#### Mouse models

Female C57BL/6J mice (Charles River Laboratories), 3 months (young) or 18-22 months (aged) old were group-housed under standard laboratory conditions with ad libitum food and water. All animal work was conducted in accordance with the UK Animals Scientific Procedures Act 1986 (PIL I44558708, and under the project licenses P8DC5B496//PP6246123 (Prof. Basson) and PA2F764C5 (Prof. Giese) and was approved by the King’s College London Animal Welfare and Ethical Review Board (AWERB).

#### ORY-1001 administration

Mice were randomly assigned to either the control (water) or ORY1001 group. Mice were given free access to drinking water containing ORY-1001 (0.27ug/mL) for 3 weeks following a 5 day on, 2 days off schedule. Behavioral or LTP studies were then conducted.

#### Cell lines

HT-22 cells were cultured at 37°C and 5% CO_2_ in DMEM, high glucose (Gibco), 10% heat-inactivated FBS (Gibco) and 1% penicillin/streptomycin (Gibco). Cells were passaged just before flasks reached 70% confluence.

### Western Blotting

Brain hippocampi were dissected and lysed to extract protein. Protein concentration was determined using a BCA kit. Samples were loaded onto a 4-15% gel and run at 150mV for 30-50min. Following transfer, membranes were blocked with 5% BSA in TBS-T for 1h at room temperature, then incubated with the primary antibody in 5% BSA in TBS-T at 4°C overnight. Membranes were incubated with anti-rabbit HRP-conjugated secondary antibody in 5% BSA in TBS-T for 1h at room temperature. Visualisation was performed using an ECL kit. The following antibodies were used: anti-H3K4me3 (CST, 9751S, 1:1000), anti-H4K12ac (Abcam, ab177793, 1:1000), anti-H3 (9715S, 1:5000), anti-tubulin (Upstate, 05-829, 1:5000), and goat anti-rabbit, and anti-mouse HRP secondary antibodies (ThermoFisher, #31460, and Proteintech, #SA00001-1, respectively, 1:5000).

### Immunofluorescence

Mouse brains were embedded in OCT and snap frozen in liquid nitrogen, Slices were cut at 25µm, washed, blocked, permeabilised, and incubated overnight with primary antibodies at 4°C. After washes, secondary antibodies were diluted in blocking buffer and incubated for 2h at room temperature. Nuclei were stained with Hoechst33342 (ThermoFisher, (1:5000). The following antibodies were used: H3K4me3 (CST, 9751S, 1:200), Arc (Synaptic Systems, 156003, 1:500), goat anti-rabbit 647 (ThermoFisher, A-21245, 1:500). Images were captured on the Zeiss Axio Observer 7 microscope and analyzed either on ImageJ or on QuPath.

### ChIPmentation and analysis

Snap-frozen hippocampi were processed using the CHIPmentation Kit for Histones, with an antibody against H3K4me3 (Diagenode). AMPure XP beads were used to clean up the libraries, which were sequenced (50bp, paired-end) on Illumina HiSeq 2000. Reads were aligned to the mouse reference genome (mm39) using Bowtie2, with an insert size of 500 bases. SAMtools was used to filter high-confidence reads. MACS3 was used to identify enriched regions (peaks), and Bedtools used to normalise read counts. limma and tidyverse (ggplot2) were used to identify differential histone mark enrichment between various conditions, defined by a p-adj value < 0.05 and a log2 fold change threshold of ±1. GenomicRanges, ChIPseeker, and rGREAT were used for annotation, and only promoter-annotated peaks taken forward for subsequent analysis.

### Behavioural Tests

Behavioural tests were conducted as previously described for OLM [28], NOR [29], and CFC [30]. Mice were handled for 2min per day for 5 days before experiments. Mice were habituated to the testing room for at least 30min before experiments. Mice were sacrificed 3h after the CFC for downstream molecular experiments.

### High-throughput Screening

All compounds were formulated at 10mM. HT-22 cells were split and seeded onto 96-well plates at a density of 20K cells/well. The Integra ViaFlo was used the day after cells were seeded for plate stamps.

HT-22 cells were split and seeded onto 96-well plates at a density of 20K cells/well in and left to proliferate for 24h. DMSO controls were split across the plate to monitor intra-plate variability. The next day, a compound intermediate dilution plate was made by thawing the OEB3 and OEB4/5 stamp plate at room temperature (Supplementary Tables 2-5). Three concentration points were tested: 10µM, 1µM, and 0.1µM. Cells were incubated with compounds 24h and subsequently fixed in 4% PFA.

For dose-response curve experiments, we used a 10-point, 3-fold titration of the compounds. Stock plates of the drugs were created to range from 1mM to 75nM and administered at a 1:10 dilution to the cells to end up with a final range of 100µM to 7.5nM.

### Immunocytochemistry

Cells were fixed with 4% PFA, after which they were washed with PBS, blocked with 5% BSA for 1h, and incubated in the primary antibody overnight at 4°C (or for 2h at room temperature), and 24h after, with the secondary antibody for 1h at room temperature. Plates were imaged on InCell Analyser 2000.

### Electrophysiology experiments

Hippocampal slices were prepared as previously described [31]. Coronal hippocampal slices (340µm) were prepared before being transferred to an interface recording containing preheated artificial cerebrospinal fluid (aCSF). Recordings began following at least 2h of incubation. Field excitatory postsynaptic potentials (fEPSPs) were recorded from CA1b stratum radiatum apical dendrites using a single glass pipette filled with 2M NaCl (2-3 MΩ) in response to orthodromic stimulation of Schaffer collateral-commissural projections in CA1c stratum radiatum. After establishing a 20min stable baseline, LTP was induced by delivering a single episode of 5 ‘theta’ bursts, each burst consisting of four pulses at 100Hz and the bursts themselves separated by 200msec (i.e., theta burst stimulation, TBS).

## Results

### H3K4me3 dysregulation in the aged mouse hippocampus

To model age-associated hippocampus-dependent memory deficits in aged mice, 24-month-old (aged) female C57BL/6J mice were compared to 3-month-old (young) mice in a standard contextual fear conditioning (CFC) test. Aged mice exhibited significantly reduced freezing 24h after an unconditioned stimulus of 33×0.7mA foot shocks, each lasting for 2s, compared to young mice (Fig. 1A). These findings recapitulate previous observations in male [4, 9, 14, 32–34] and female [5, 10, 35–36] mice.

**Fig. 1.**
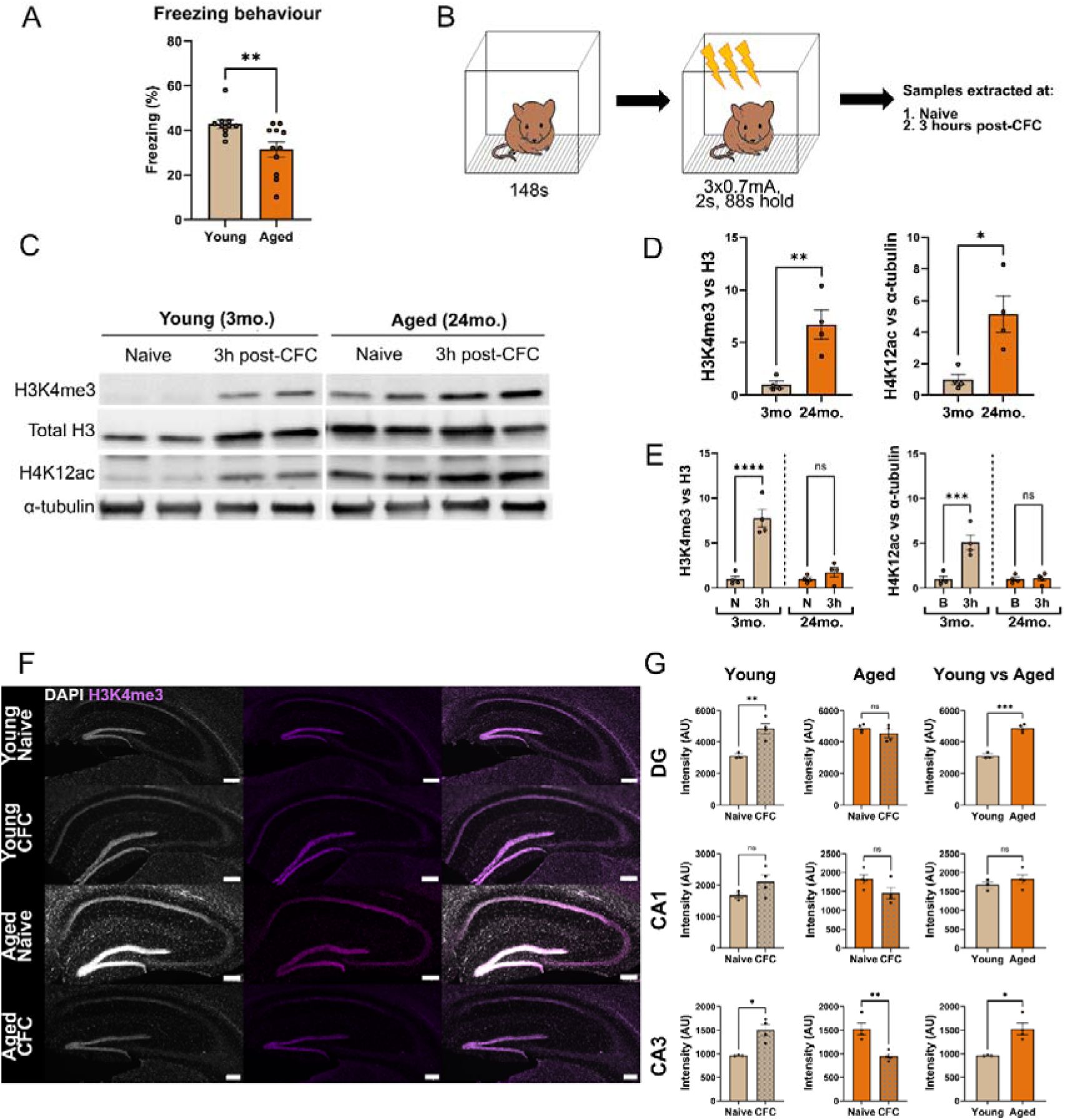
CFC-dependent H3K4me3 induction is impaired in the hippocampus of aged mice. (A) Results showing freezing behaviour, an indication of associative memory, in young and aged mice 24h after CFC (n=10/11 per condition). (B) Schematic showing the CFC protocol for molecular experiments. (C) Representative images of Western blots for H3K4me3, total H3, H4K12ac and a-tubulin on total cell lysates from young and aged hippocampi (n=2 shown, repeated 3 times with 4 independent samples showing the same results). (D) Quantification of signal intensity of H3K4me3 levels relative to total H3 and H4K12ac levels relative to α-tubulin (n=4 young (3mo.) and old (24mo.) mice). (E) Relative quantification of H3K4me3 and H4K12ac in hippocampi from young (3mo.) and aged (24mo.), naive (N) and CFC mice (3h) (n=3/4 mice per condition). Two-way ANOVA was conducted, followed by Sidak’s post-hoc tests. (F) Images of the hippocampi of young and aged female mice, stained for H3K4me3 (purple) and DAPI (grey). Scalebars represent 200µm. (G) Analysis of H3K4me3 intensity in different regions of the hippocampus, with comparisons shown for young and aged mice, naive or 3h after CFC, across DG, CA1, and CA3 regions. Each dot is a biological replicate. n=3/5 slices imaged each from n=3/4 mice per condition. Student’s t-tests were conducted. Data are presented as mean ± SEM. * p<0.05, ** p<0.01, *** p<0.001, **** p<0.0001.

To determine if H3K4me3 is activity-regulated in the mouse hippocampus, we first performed a short time course and found that H3K4me3 induction was evident 3h after CFC (Fig. S1A, B). Next, we compared hippocampi from young and aged female mice 3h after CFC with hippocampi from naive age-matched mice (Fig. 1B). Western blot analyses revealed significantly elevated levels of H3K4me3 in hippocampi from naive, aged mice compared to naive, young mice (Fig. 1C, D). H4K12ac was similarly elevated in the hippocampus of naive, aged mice relative to naive, young mice, consistent with results from Peleg et al. (14, Fig. 1C, D). An activity-induced increase in H3K4me3 was observed in young, but not aged mice in response to CFC. (Fig. 1C, E; F(1,12) = 24.42, age x CFC effect p=0.0003). The activity-induced H4K12ac observed in young mice was also absent in aged mice, as previously reported (Fig. 1E; F(1,12)=16.80, age x CFC effect p=0.0015 [14]

To confirm these observations, H3K4me3 was visualised in the hippocampus by immunostaining (Fig. 1F). Significant H3K4me3 induction was visible in the dentate gyrus (DG) and area CA3 in young mice after CFC with a similar trend in area CA1 (Fig. 1G). No induction was observed in aged mice, with H3K4me3 levels declining in area CA3 after CFC (Fig. 1G). H3K4me3 levels were significantly higher in naïve aged mice compared to young controls in the DG and area CA3 (Fig. 1G).

To understand how these H3K4me3 differences relate to the induction of proteins involved in synaptic plasticity and memory formation, hippocampal sections were immunostained for Arc (IEG, Fig. S1C). Young mice showed an increase in Arc+ cells in all regions after CFC (Fig. S1D), while aged mice had few Arc+ cells and no induction after CFC (Fig. S1D).

### H3K4me3 alterations around promoter regions in the aged hippocampus

H3K4me3 is typically enriched around TSSs of actively transcribed genes. Using ChIPmentation, H3K4me3 distribution around TSSs in hippocampal tissue was visualised [37]. Comparing H3K4me3 enrichment around TSSs of naive, young and aged mice revealed a net increase in H3K4me3 signal in old age (Fig. 2A), consistent with Western blot and immunostaining data (Fig. 1) Differential peak calling identified more genes with increased than decreased H3K4me3 at TSSs (370 increased and 315 decreased, Fig. 2B). Gene ontology (GO) enrichment analysis revealed small GTPase mediated signal transduction and cytoskeletal organisation as the most significant, whilst also including chromatin remodelling, a known feature of aged neurons [38] and post-synaptic specialisation organisation (Fig. S2A).

**Fig. 2.**
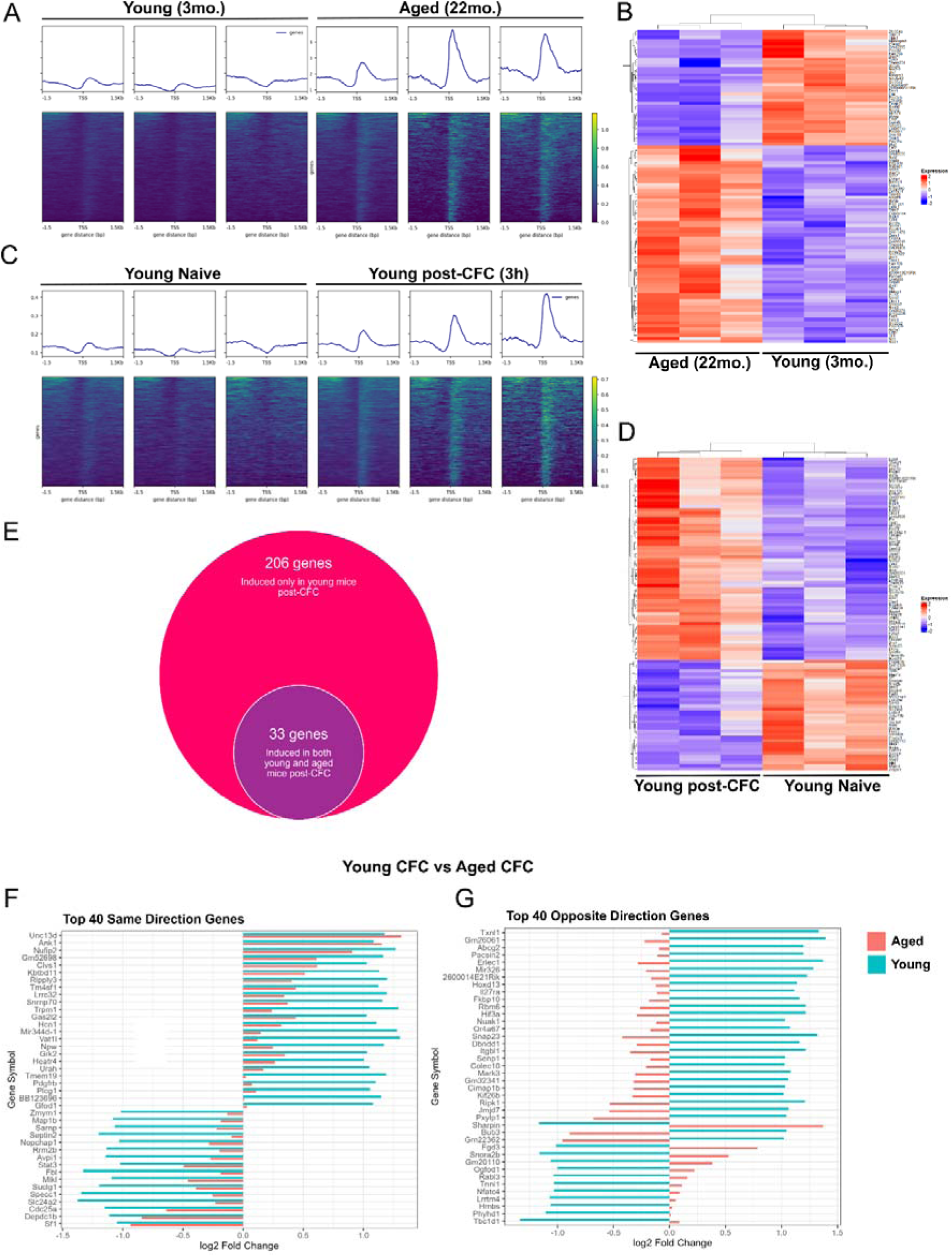
Genome-wide age-associated changes to H3K4me3. (A) Normalised H3K4me3 peaks at TSSs in young and aged hippocampi (n=3 mice for each age group). Heatmaps accompanying each peak comparison showing TSSs with altered H3K4me3 peaks in aged mice. (B) Heatmaps showing the top 100 upregulated and downregulated H3K4me3 TSSs between young and aged mice. (C) Normalised H3K4me3 peaks at TSSs in young hippocampi from naive mice and 3h after CFC (n=3 mice for each group). Heatmaps accompanying each peak comparison showing TSSs with altered H3K4me3 peaks induced by CFC. (D) Heatmaps showing the top 100 upregulated and downregulated H3K4me3 TSS peaks induced by CFC in young mice. (E) Venn diagram showing number of overlapping differentially enriched genes in an interaction model comparing young and aged mice, before and after CFC. (F, G) Bar plots showing changes in the magnitude in log2 (fold change) values between young and aged mice post-CFC of top 40 genes around which H3K4me3 enrichment changes in the same direction (F) and different directions (G). Note the blunted response for most genes in aged mice in F and genes with the opposite effect in G.

Next, H3K4me3 peaks were compared in hippocampi obtained from naive, young mice and young mice 3h after CFC. Genes with clear increases in H3K4me3 at their TSSs in response to CFC were identified (Fig. 2C, D), while some showed the opposite (Fig. 2D). GO terms included actin filament organisation, neurogenesis, metabolism, and cytoskeletal rearrangement (Fig. 2D, S2B). We repeated this analysis with hippocampi from aged mice. Relative to their naive controls, aged mice showed highly variable H3K4me3 enrichment around TSSs after CFC (Fig. S2C). Genes with significant differences in H3K4me3 after CFC (Fig. S2D), were categorised as being involved in neurogenesis, Wnt signalling, gliogenesis, and dendrite development (Fig. S2E).

Of the 239 genes that showed a significant induction of H3K4me3 levels at the TSSs after CFC in young mice, only 33 genes showed a significant induction in aged mice (Fig. 2E). The remaining 206 genes implicated in activity-dependent induction were unaltered in aged mice, consistent with a deficit in activity-induced H3K4me3 (Supplementary Table 1). GO term analysis of these 206 genes revealed a significant enrichment of genes associated with synaptic plasticity, synapse maturation, neurogenesis, and synaptic vesicle processing. Analysis of the 33 genes conserved across ages revealed H3K4me3 enrichment around genes associated with protein catabolism and cell division (Fig. S2F). A comparison of H3K4me3 levels at genes with most significant H3K4me3 enrichment after CFC, revealed genes where H3K4me3 changed in the same direction (increased/reduced) in young and aged hippocampi, (Fig. 2F), and genes where H3K4me3 changed in the opposite direction (Fig. 2G). In the former group, the activity-induced H3K4me3 change was weaker at most genes in aged mice (Fig. 2F). The latter group suggests that the activity-induced mechanisms that mediate H3K4me3 changes at TSSs in young mice were not only weaker, but also qualitatively different in aged mice (Fig. 2G). These results demonstrate significantly altered H3K4me3 enrichment around TSSs in the ageing hippocampus.

### High throughput screening to identify H3K4me3-altering compounds

To determine if targeting H3K4me3 regulators in vivo could restore learning and memory in aged mice, we first developed a high throughput, immunocytochemistry-based assay in the immortalised hippocampal cell line, HT-22 [39]. To identify a suitable positive control that can robustly alter H3K4me3 levels, we tested compounds previously shown to influence learning behaviour in *in vivo* experiments. HT-22 cells were cultured for 24h in the presence of WDR5-0103 [40] GSK-LSD1 [41] or AR-42 [42] in 10-point dose response curves, followed by fixation and H3K4me3 immunostaining (Fig. 3A). Neither WDR5-0103, nor GSK-LSD1 had a significant effect on H3K4me3 levels in HT22 cells (Fig. 3A), but AR-42 increased H3K4me3 levels in a concentration-dependent manner (Fig. 3A). Western blots confirmed the dose-dependent induction of H3K4me3 in HT-22 cells upon AR-42 administration (Fig. 3B, 3C; F(2, 15)=13.16, p=0.0005). Therefore, AR-42 was selected as the positive control for the ICC-based assay.

**Fig. 3.**
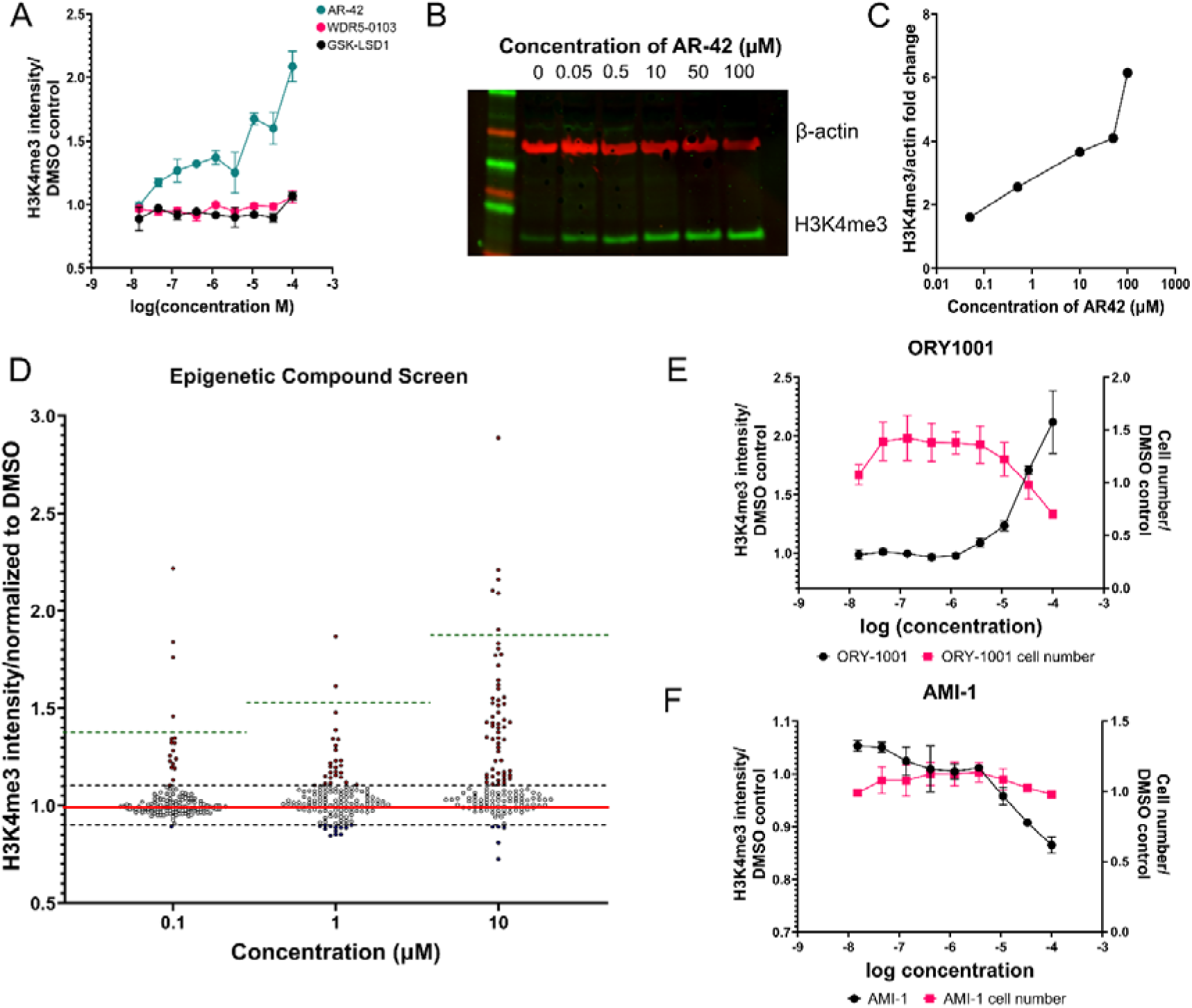
High throughput screening identifies ORY-1001 as robust regulator of H3K4me3. (A) Dose-response curves showing H3K4me3 intensity in response to AR-42, GSK-LSD1 or WDR5-0103 in an ICC-based assay in HT-22 cells. (B) Western blot comparing H3K4me3 and b-actin protein levels in HT22 cells, treated with increasing concentrations of AR-42 for 24 hours to validate B. (C) Quantification of H3K4me3 levels in HT22 cells, relative to β-actin, from Western blots. (D) Results of the epigenetic compound screen. Each individual point is a different compound. Compounds were tested at 0.1, 1, and 10µM. H3K4me3 intensity upon AR-42 treatment as a positive control was plotted on the graph as a dotted green line. Grey dotted lines represent a 10% increase or decrease in H3K4me3 intensity, as an arbitrary threshold for active compounds. Red dots show compounds with greater than 10% increase in H3K4me3 intensity while blue dots show compounds with greater than 10% decrease in H3K4me3 intensity. (E) Dose-response curve of ORY-1001, lead H3K4me3 stimulating compound identified in the screen. (F) Dose-response curve of AMI-1, lead inhibitor of H3K4me3 identified in the screen. Data are presented as mean ± SEM. n=3 for dose-response curve experiments.

The assay was optimised to account for cell-seeding density (Fig. S3A) and duration of treatment (Fig. S3B). H3K4me3 immunocytochemistry showed the most robust increase after a 24h treatment with AR-42, in seeding density of 20,000 cells in 96-wells. The assay was tested for replicability and robustness, evidenced by a Z’-score of 0.43 for 24h of treatment, and a signal-to-noise ratio of 4.11, indicative of a sensitive assay, with true signals being easily distinguishable from background (Fig. S3C).

A library of 145 epigenetic compounds targeting methyltransferases, demethylases, acetyltransferases, deacetylases, kinases, chromatin remodelling complexes, epigenetic reader and writer domains was screened at 0.1µM, 1µM, and 10µM (Supplementary Tables 2-4, Fig. 3D). In total, 54 compounds modulated H3K4me3 levels in HT-22 cells, with 42 increasing (Fig. S4A) and 12 reducing H3K4me3 levels (Fig. S4B). Among these, compounds with the greatest changes in H3K4me3 levels were explored further. Most of these compounds were eliminated due to their toxicity profiles (Fig. S4C). The LSD1 (KDM1A) lysine demethylase inhibitor ORY-1001, showed a dose-dependent increase in H3K4me3 levels with minimal toxicity (Fig. 3E). AMI-1, an arginine methyltransferase inhibitor, showed a dose-dependent decrease in H3K4me3 without causing significant cell death (Fig. 3F). Only ORY-1001, among several LSD1 inhibitors tested, increased H3K4me3 levels while maintaining cell viability, and was chosen as the candidate compound for further experiments.

### ORY-1001 rescues learning and memory in aged mice

ORY-1001 can be administered in drinking water and cross the blood-brain barrier [41]. ORY-1001 was administered to 22-month-old mice in drinking water (0.02mg/kg in water, 5 days on, 2 days off), 2 weeks prior to and during the behavioural experiments. We first assessed hippocampus-dependent memory using the object location memory task (OLM) (Fig. 4A). All mice explored both locations equally in the training session (Fig. 4B). In the test session 24h after, young mice spent significantly more time exploring the object in a novel location, compared to the object in the original location, whilst aged mice failed to discriminate between these objects, indicating spatial memory impairment (Fig. 4C). ORY-1001 rescued spatial learning and memory, with ORY-1001-treated aged mice having discrimination indices (DI) comparable to young mice (Fig. 4C, F(1,46)=6.32, age x drug effect p = 0.016).

**Fig. 4.**
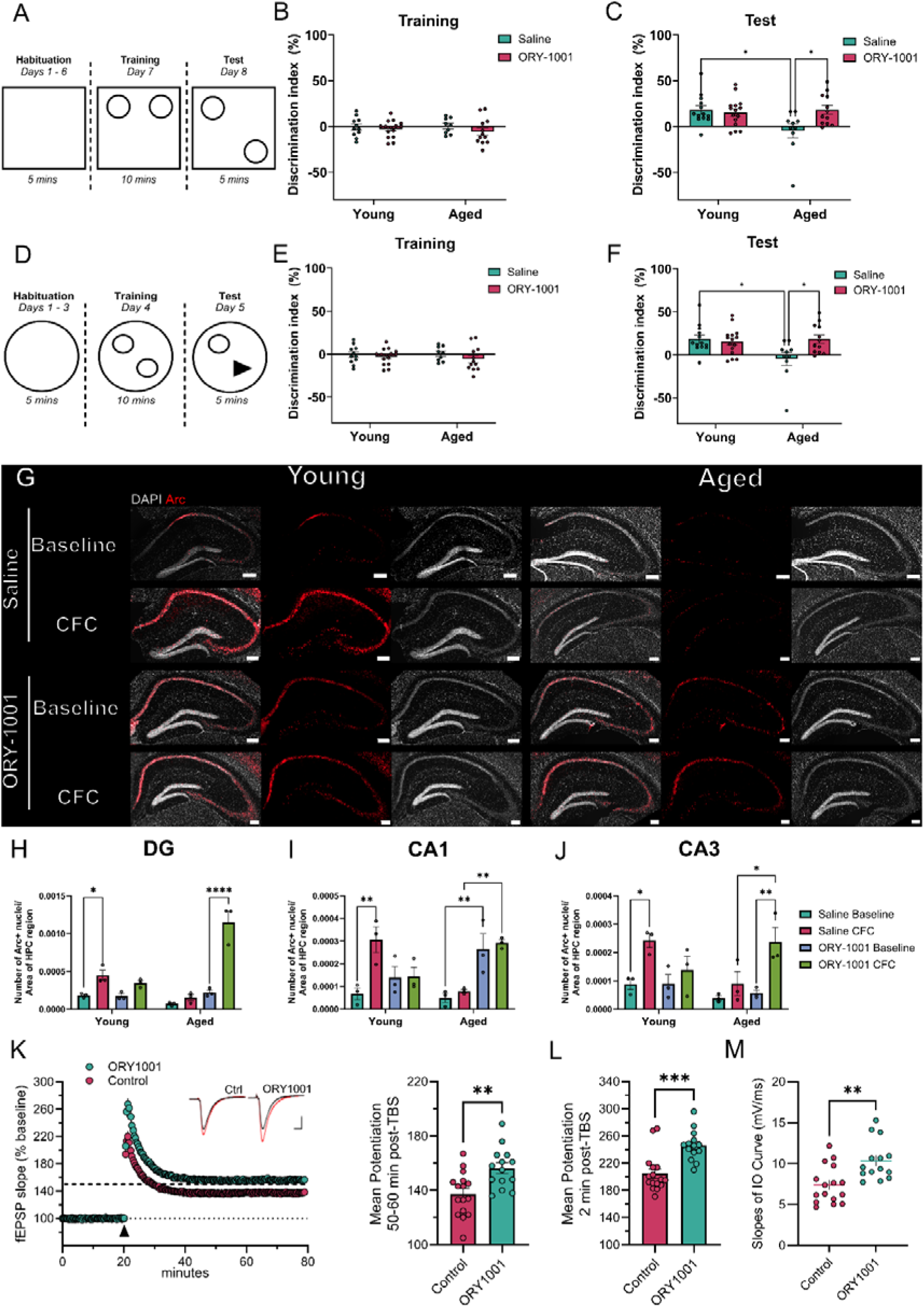
ORY-1001 treatment restores learning and memory in aged mice. (A – C) OLM task. (A) Schematic showing the experimental design of the OLM task. (B) Discrimination index (DI) during the Training session, measured by comparing time spent exploration both objects. (C) DI comparing the novel object location and the original object location in the test session. (D – F) NOR task. (D) Schematic showing the experimental design of the NOR task. (E) Discrimination index (DI) during the training session, measured by comparing time spent exploration both objects. (F) DI comparing the novel object and the original object. (G) Representative images of young and aged mice at baseline or post-CFC, given either ORY-1001 or saline, stained with DAPI (grey) or Arc (red). Scalebars = 200µm. (H-J) Quantification of Arc+ nuclei normalised by area of hippocampal region: Dentate Gyrus (DG, H), CA1 (I) and CA3 (J). Each dot represents a biological replicate, n=3 mice per condition, n=3-5 slices per animal. Note the increase in Arc+ nuclei upon CFC in young mice, the blunting of this response in aged mice and the significant increase in Arc+ nuclei in aged mice upon ORY-1001 treatment. (K) Left, fEPSP slope recordings in hippocampal slices from aged mice given either saline or ORY-1001. The dotted line indicates the average LTP in young mice measured over 5 years in the Wood laboratory (Keiser et al., 2021; Kwapis et al., 2018; Tran et al., 2023; Vogel-Ciernia et al., 2013). Scale bar: 1mV per 5ms. Right, Summary graph showing mean fEPSP slope 50-60m after stimulation. (L) Summary graph showing mean fEPSP slow 2m after stimulation to highlight short-term LTP. (M) Summary graph showing individual slopes of the input/output curve collected from a slice in each group. N=13/16 slices per condition used. 3mo. mice n=26, 20mo. mice n=26 for behavioural data. Data are presented as mean ± SEM. Two-way ANOVA was conducted in C, F, H-J, followed by Sidak’s post-hoc tests. Welch’s t-tests were conducted for K - M. * p<0.05, ** p<0.01, *** p<0.001, **** p<0.0001.

Learning and memory were also assessed in the novel object recognition (NOR) task (Fig. 4D). NOR involves both the hippocampus and perirhinal cortex, with the perirhinal cortex representing basic information about familiarity or novelty of an object, with the hippocampus required for object memorisation by encoding information about the experience of the object [43]. All mice explored the objects equally in the training session (Fig. 4E). Aged mice administered saline showed memory impairment in the test session, which was restored by ORY-1001 administration (Fig. 4F, F(1,46)=7.85, age x drug interaction effect p<0.0074, age effect p=0.017).

To assess the effectiveness of ORY-1001 in restoring activity-dependent Arc induction after CFC, mice were administered ORY-1001 intraperitoneal for 3 days before the CFC task [41]. Young mice showed an increase in Arc+ cells in all hippocampal areas post-CFC, while aged mice did not (Fig. 4G, H, I, J). However, aged mice treated with ORY-1001 showed a striking activity-dependent increase in Arc+ cells in the DG (Fig. 4H; F(1, 16) = 27.93, age x drug x CFC effect p<0.0001), and CA3 (Fig. 4J; F(1,16)=6.09, age x drug x CFC effect p=0.025), and a similar increase in Arc+ cells in the CA1 of naive aged mice compared to vehicle-treated mice (Fig. 4I; F(1,16)=20.71, age x drug effect p=0.0003).

LTP, a form of synaptic strengthening thought to underlie learning, can be studied in ex vivo brain slices [12, 44–45]. Acute hippocampal slices were used to evaluate changes in synaptic plasticity in area CA1 from aged mice. ORY-1001 treatment led to a significant increase in both short-term and long-term potentiation in aged hippocampi, assessed by the mean potentiation at 2min and 50-60min post-TBS, respectively (Fig. 4K, L). The observed marked shift in slope of the input/output (IO) curve in slices from ORY-1001-treated mice relative to controls (Fig. 4M), suggests increased neuronal excitability in the hippocampus. Together, this data shows that ORY-1001 treatment can reverse learning and memory deficits, restore activity-dependent Arc induction and increase LTP in aged mice.

## Discussion

In this manuscript, we report significant age-associated changes in H3K4me3 in the hippocampus of aged female mice. We found a significant increase in H3K4me3 levels in naive mice, but the induction of H3K4me3 in the hippocampus after CFC is markedly blunted in aged mice. Genome-wide ChIP analysis identified genes that undergo activity-dependent H3K4me3 changes at their TSSs in young mice and most of these showed either a blunted or different response in aged mice. Using a high-throughput immunocytochemistry-based screen, we identify ORY-1001, a histone demethylase inhibitor, as a suitable compound for targeting the H3K4me3 machinery in vivo. ORY-1001 treatment rescued spatial and novel object recognition memory and LTP in aged mice.

Previous studies have identified several histone PTMs that are regulated by neuronal activity in the hippocampus during learning and memory consolidation, including H4K12ac, H3K9me2, H3K27me3, and H3K4me3 [21, 32, 46–47]. In this report, we used independent experimental approaches to show that H3K4me3 is increased in the aged hippocampus, that H3K4me3 is induced by activity in the young hippocampus and that this induction is significantly blunted and changed in the aged hippocampus. Previous studies have also reported increased H3K4me3 levels in the prefrontal cortex of Alzheimer’s disease models [13, 40] and age-related increases in H3K4me2 at stress-response genes in rodent and macaque brains [48–49], suggesting that H3K4me changes may underlie memory deficits in neurodegenerative conditions. These increases in H3K4me2/3 may indicate a general shift towards a more permissive chromatin environment, as indeed suggested by reports of global DNA hypomethylation in the aged hippocampus [50–53].

Genome-wide analysis of H3K4me3 differences between young and aged mice implicate genes involved in GTPase-mediated signal transduction, actin filament organisation, and chromatin remodelling. Altered GTPase activity associated with endocytosis and autophagy, and actin dynamics crucial for LTP [54][55–57] are linked to memory stabilisation and rely on the IEG, Arc [57–59]. Age-associated deficits in activity-induced H3K4me3 implicate genes involved in neurogenesis, dendrite development, and synaptic maturity, essential for long-term memory [60–62]. Genome-wide analysis of activity-induced H3K4me3 changes suggests that some of these changes were not merely blunted but changed in the opposite direction in aged mice, suggesting that different mediators are recruited to H3K4me3 upon neuronal activation. Further work is needed to test these hypotheses and identify the mechanisms responsible.

To therapeutically reverse aberrant H3K4me3 levels in the ageing hippocampus, we developed an immunocytochemistry-based high-throughput screening model. Out of the 145 screened compounds, 54 altered H3K4me3 levels in vitro. Most identified compounds were HDAC inhibitors, highlighting the interconnectedness of histone modifications and their potential crosstalk [63–64]. ORY-1001 was chosen for in vivo experiments and restored normal learning and memory in the OLM and NOR tasks. Interestingly, ORY-2001, a related compound, has been shown to increase IEG expression and rescue NOR deficits in an accelerated ageing model [65].

Activity-dependent Arc induction in young mice [66–69], and its subsequent impairment in the aged hippocampus have been studied previously [70–72]. We demonstrated that ORY-1001 can restore activity-dependent Arc induction in aged mice, especially in the DG and CA3 regions. Aged mice also have impaired LTP [73–75], and ORY-1001 can increase LTP to levels similar to young mice [31, 76–77]. The increases observed in the I/O curve and short-term potentiation suggest that ORY-1001 may contribute towards altered neurotransmitter activity or neuronal excitability in aged mice, influencing LTP [78]. This increased excitability can be explained by the elevated Arc levels in the CA1 region of aged mice given ORY-1001, as Arc regulates intrinsic excitability of rat hippocampal neurons [79]

These results are also consistent with the increased recruitment of KDM1A to transcription factors that modulate IEGs upon administration of related compound ORY-2001 [65]. However, LSD1 inhibition affects many processes, so more work is needed to understand the mechanisms whereby ORY-1001 restores learning and memory in aged mice.

We believe that ORY-1001 treatment restores learning and memory primarily by restoring activity-induced gene regulation. However, ORY-2001 can reduce the levels of inflammatory cytokines [80], so these anti-inflammatory mechanisms may also contribute to our findings. We acknowledge the use of predominantly single sex mice as a limitation. We confirmed restoration of activity-dependent Arc induction in male mice, but further work is needed to evaluate ORY-1001 in aged male mice. Together, our results show that ageing disrupts H3K4me3 levels, and that targeting this PTM with a histone demethylase inhibitor improves learning and memory.

## Supporting information

ChIP-Seq Data

## Acknowledgments

We thank Dr. Jordi Senserrich Velasco (ORYZON) for advice on ORY-1001 preparation and administration, the BSU staff at King’s College London and University of Exeter for supporting animal experiments, and Lily Meyer for helping with behavioural data analysis. All animal experiments were approved by institutional AWERBs and were performed under Home Office Project licence PP6246123. This work was supported by the MRC DTP iCASE programme, King’s College London and MSD, to SUB, the Wellcome Trust NIIHD PhD programme, King’s College London to AG, National Institute for Health and Care Research & the Medical Research Council grant MR/Y008170/1 to MAB, Medical Research Council grant MR/V013173/1 to MAB and KPG, NIH/NIA R01 grant AG076835 to MAW and start-up funds from the Faculty of Health and Life Sciences, University of Exeter to MAB.

## Author contributions

MAB conceived the project with SG and KPG. SUB designed and performed the experiments and analysed the data with contributions from LP-S, PS, JLR and AG. LM analysed the ChIPmentation data, EK and TF performed and analysed the LTP experiments. MAW, SG, KPG and MAB supervised the work. SUB and MAB wrote the manuscript with contributions from the other co-authors.

## Resource availability

### Lead contact

Further information and requests for resources and reagents should be directed to, and will be fulfilled by the lead contact, Albert Basson

### Data and code availability

Data will be uploaded to GEO and made freely available upon publication. Bioinformatics code is available at: https://github.com/lm690/Project_11657.git

## Ethics Declarations

### Declaration of generative AI and AI-assisted technologies

None.

