## Supplementary Data for "Inhibition of histone lysine demethylase restores learning and memory in aged mice"

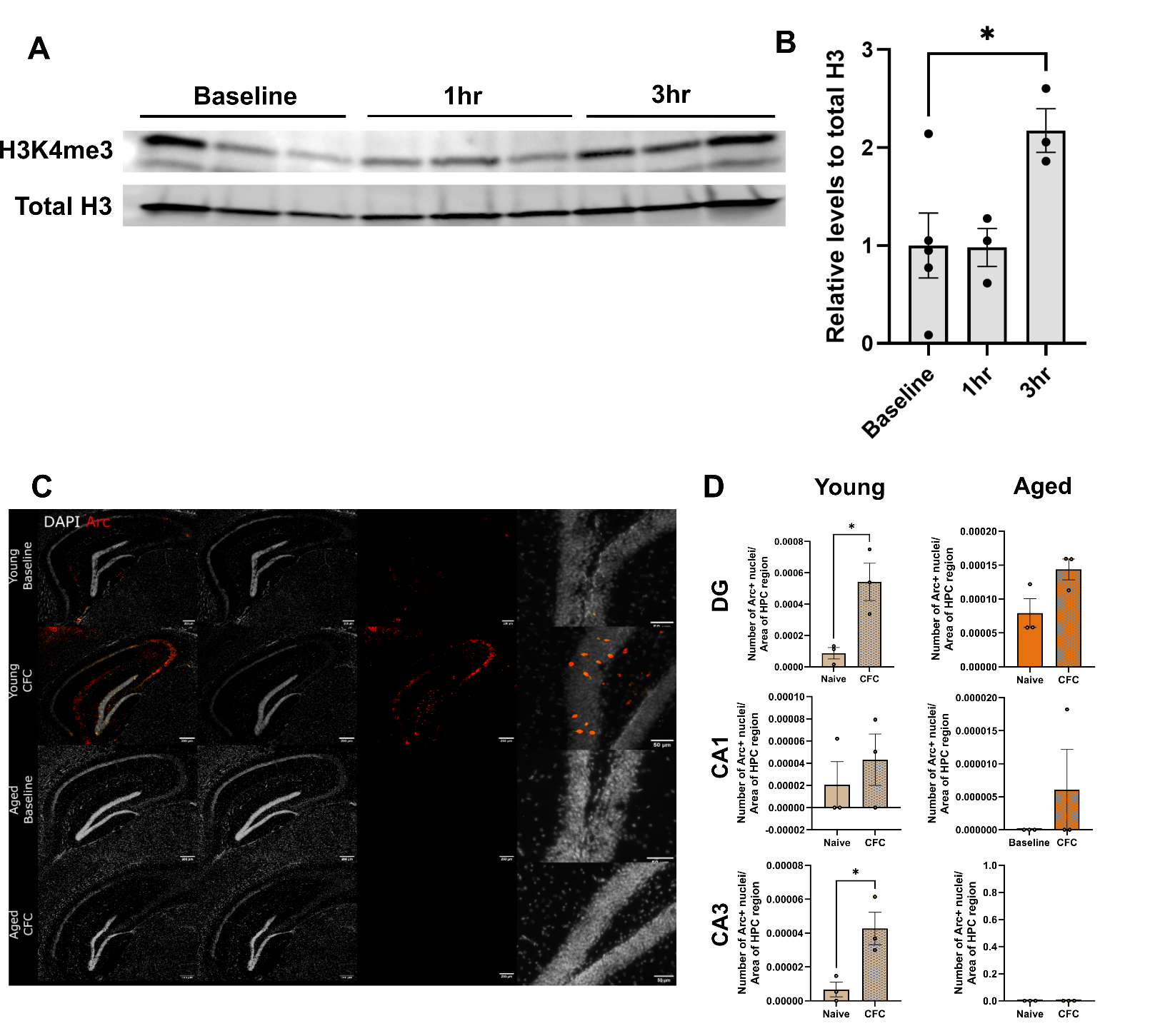

**Fig. S1. Kinetics of H3K4me3 regulation after CFC in young mice and downstream Arc expression.** (A) Representative Western blot for time-course experiment conducted on young mouse hippocampi collected at baseline, 1h, and 3h post-CFC. (B) Quantification of data in A, H3K4me3 levels normalised relative to total H3. (C) Representative immunofluorescence images of young and aged hippocampi at baseline or post-CFC, given either ORY-1001 or saline, stained with DAPI (grey) or Arc (red). Scalebars show 200um. (D) Quantifications of Arc+ nuclei normalised per area of hippocampal region. Each dot represents a biological replicate. N=3 mice used per condition. Data are presented as mean ± SEM. Data was analysed via Student’s t-test.

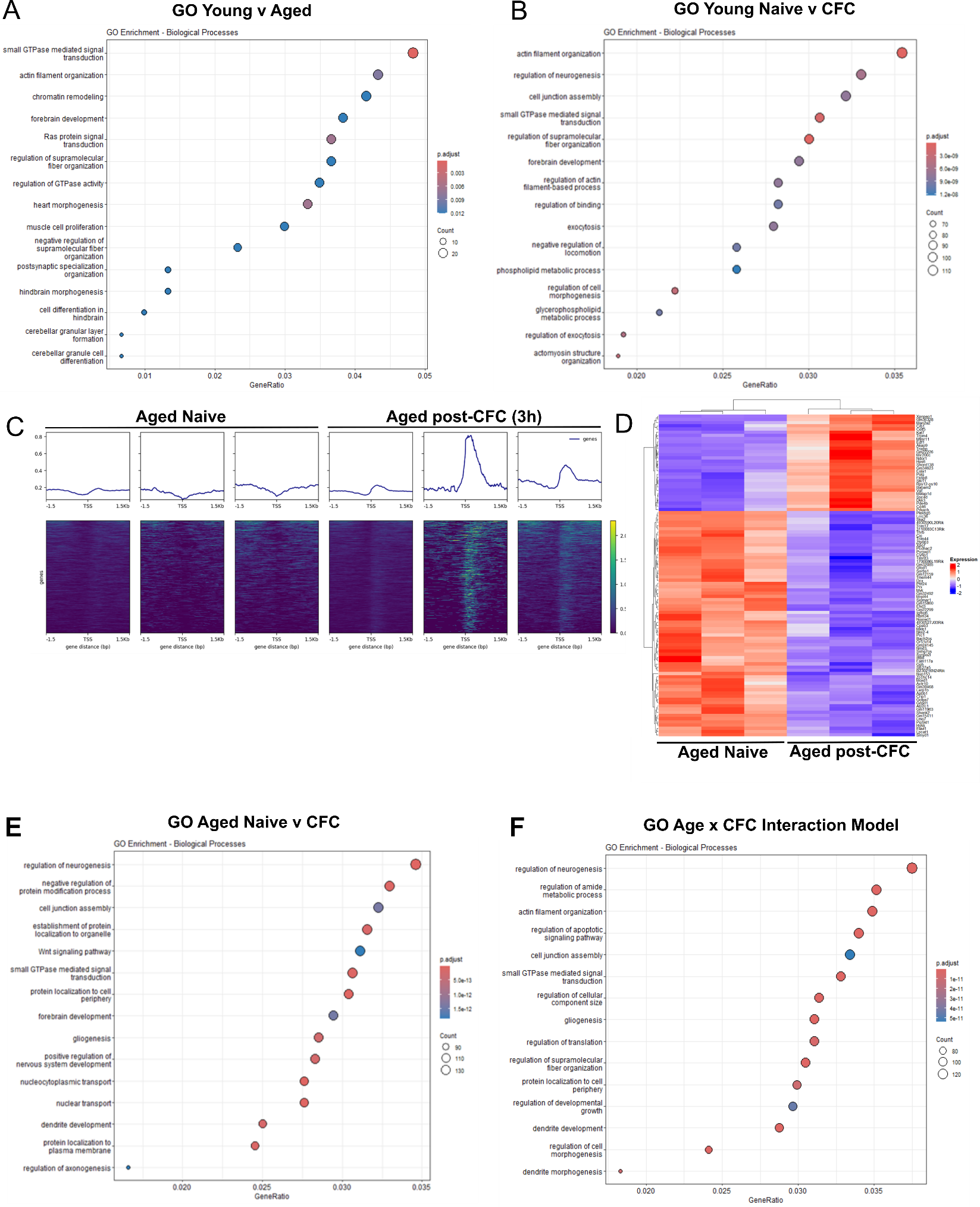

**Fig. S2. H3K4me3 enrichment is altered around TSSs in aged mice after CFC.** (A) Gene ontology enrichment maps showing altered molecular pathways with age. (B) Gene ontology enrichment maps showing altered molecular pathways with CFC in young mice. (C) Normalised H3K4me3 peaks at TSSs in naive aged hippocampi, compared to aged hippocampi 3h post-CFC (n=3 mice for each group). Heatmaps accompanying each peak comparison showing TSSs with altered H3K4me3 peaks. (D) Heatmaps showing the top 100 upregulated and downregulated H3K4me3 TSSs between aged before and after CFC. (E) Gene ontology enrichment maps showing altered molecular pathways with CFC in aged mice. (F) Gene ontology enrichment maps showing altered molecular pathways between young and aged mice, before and after CFC.

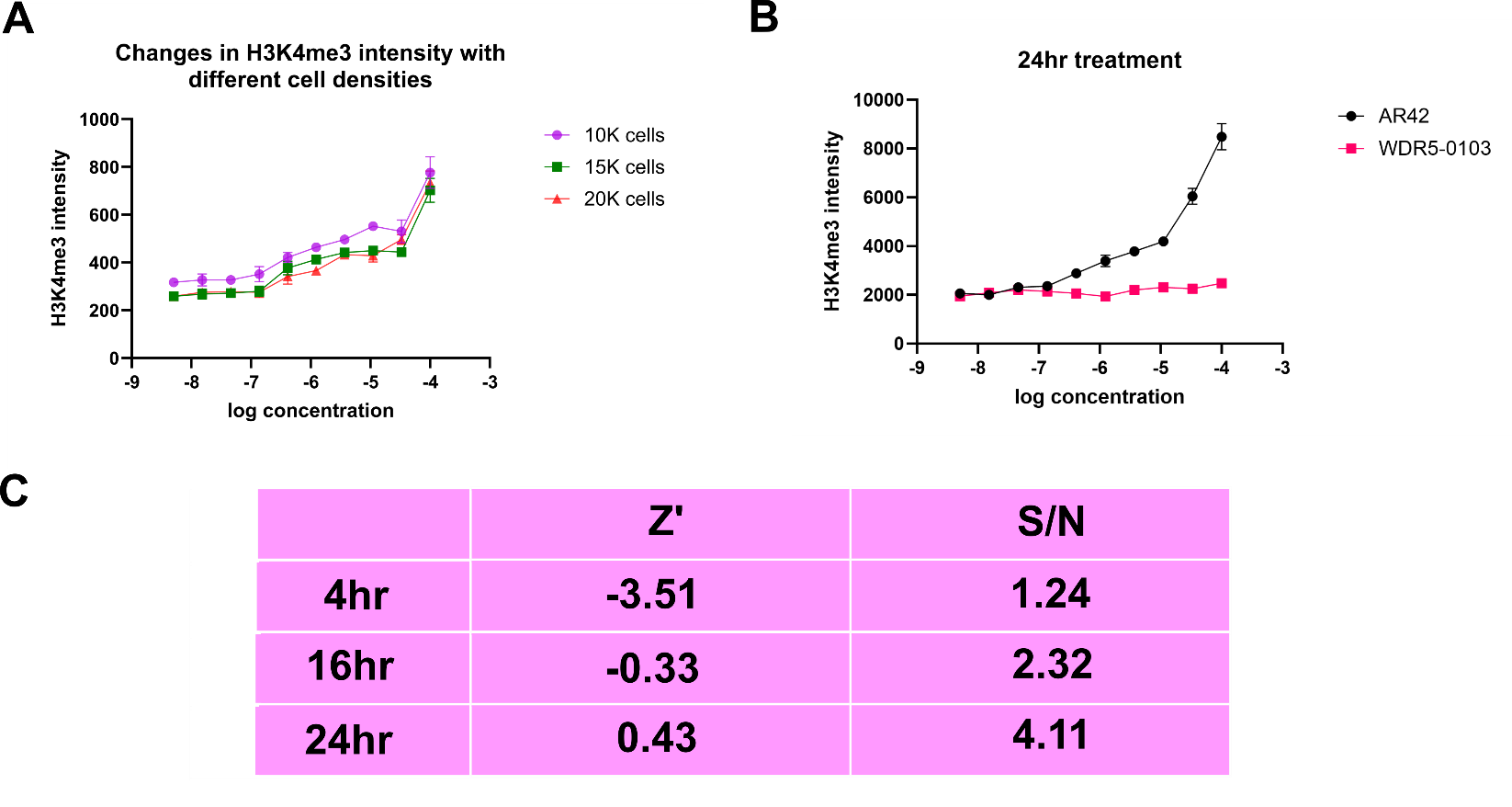

**Fig. S3. Optimisation of the ICC-based HTS assay.** (A) Dose-response curves showing changes in H3K4me3 intensity with different cell-seeding densities. (B) Dose-response curve showing changes in H3K4me3 intensity after 24h of treatment with AR-42, but not WDR5-0103. (C) Table showing Z’ values and signal/noise ratios for drug treatments of 4h, 16h, and 24h.

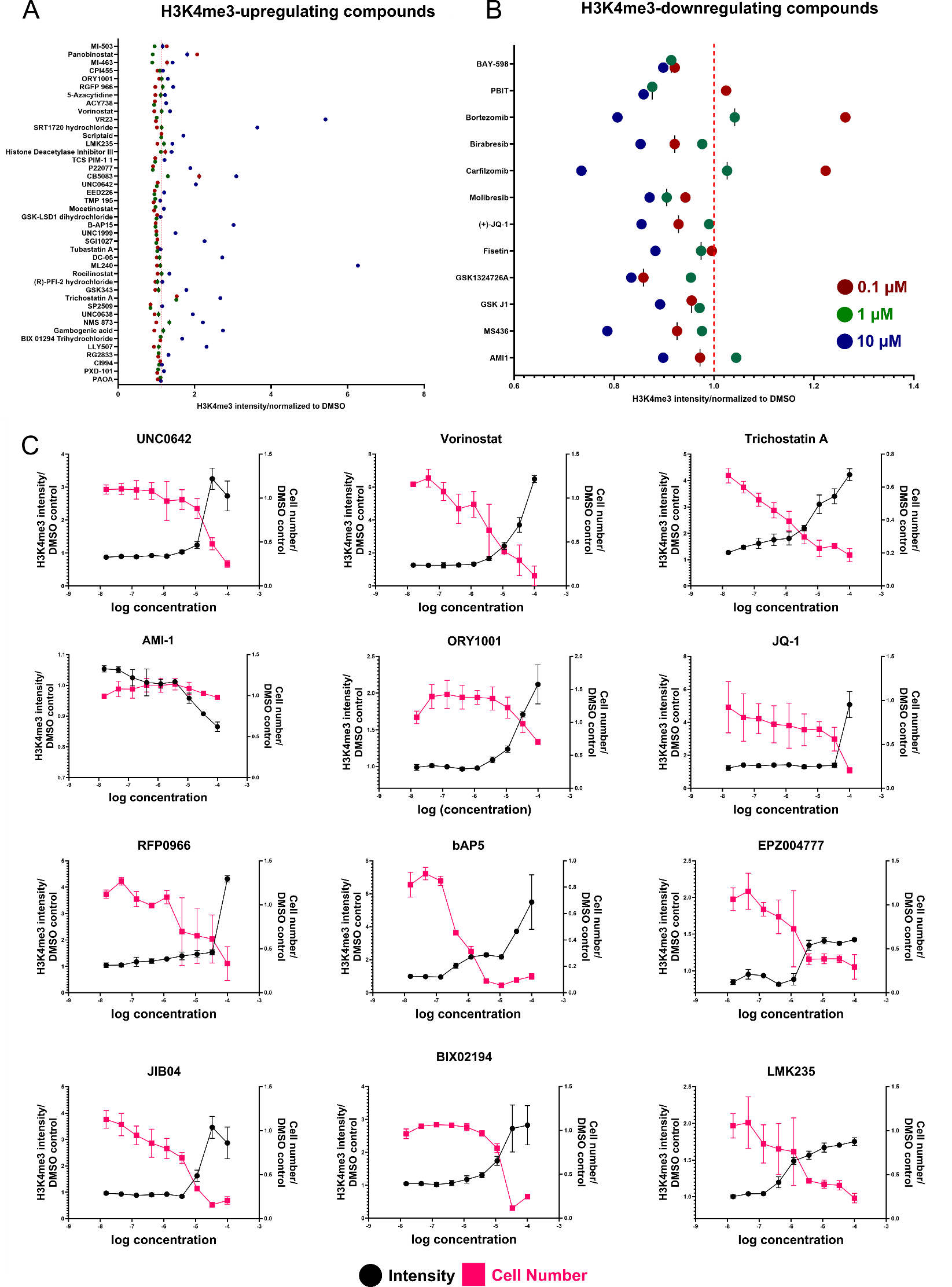

**Fig. S4. Results of the HTS assay.** (A) Figure showing compounds capable of increasing H3K4me3 levels at 0.1µM, 1µM, and 10µM. (B) Figure showing compounds capable of downregulating H3K4me3 levels at 0.1µM, 1µM, and 10µM. (C) Dose-response curves of selected compounds. (Left to right, top to bottom). UNC0642, Vorinostat, Trichostatin A, AMI-1, ORY-1001, JQ-1, RFP0966, bAP5, EPZ004777, JIB04, BIX02194, LMK235.

**Supplementary Table 1 (IN SEPARATE FILE)**

**Supplementary Table 2: Plate views of compounds from MSD compound library (3 plates used for compound screening).**

| **Empty** | **Empty** | **Empty** | **Empty** | **Empty** | **Empty** | **Empty** | **Empty** | **Empty** | **Empty** | **Empty** | **Empty** |
| --- | --- | --- | --- | --- | --- | --- | --- | --- | --- | --- | --- |
| **Empty** | **DMSO** | **AS8351** | **CI994** | **Curcumin** | **Valproic Acid** | **Piribedil** | **Anacardic acid** | **SRT1720 hydrochloride** | **Inauhzin** | **AR42** | **Empty** |
| **Empty** | **DMSO** | **Parthenolide** | **AMI1** | **NSC 3852** | **Daminozide** | **Diflunisal** | **Trichostatin A** | **Apabetalone** | **P005091** | **AR42** | **Empty** |
| **Empty** | **DMSO** | **Sirtinol** | **Benzenebutyric acid** | **Methyl L-histidinate dihydrochloride** | **Fisetin** | **Cilastatin** | **PXD-101** | **Droxinostat** | **C7280948** | **AR42** | **Empty** |
| **Empty** | **AR42** | **Scriptaid** | **Benzenebutyric acid** | **Sinapinic Acid** | **lutidinic acid** | **RG108** | **BML210** | **DBEQ** | **BCI121** | **DMSO** | **Empty** |
| **Empty** | **AR42** | **AOB2796** | **Nicotinamide** | **Theophylline** | **PR619** | **Thioguanine** | **Panobinostat** | **OF1** | **Decitabine** | **DMSO** | **Empty** |
| **Empty** | **AR42** | **ITSA1** | **Entinostat** | **5-Methyl-2'-deoxycytidine** | **Theophylline-7-acetic acid** | **1-Naphthohydroxamic acid** | **Mocetinostat** | **DC-05** | **C646** | **DMSO** | **Empty** |
| **Empty** | **Empty** | **Empty** | **Empty** | **Empty** | **Empty** | **Empty** | **Empty** | **Empty** | **Empty** | **Empty** | **Empty** |

| **Empty** | **Empty** | **Empty** | **Empty** | **Empty** | **Empty** | **Empty** | **Empty** | **Empty** | **Empty** | **Empty** | **Empty** |
| --- | --- | --- | --- | --- | --- | --- | --- | --- | --- | --- | --- |
| **Empty** | **DMSO** | **Rocilinostat** | **SGC-CBP30** | **SP2509** | **GSK1324726A** | **EPZ015666** | **UNC 926 hydrochloride** | **MS023** | **UAMC 00039 dihydrochloride** | **AR42** | **Empty** |
| **Empty** | **DMSO** | **PFI1** | **GSK343** | **GSK-LSD1 dihydrochloride** | **LLY507** | **PFI4** | **BI9564** | **EZM 2302** | **B-AP15** | **AR42** | **Empty** |
| **Empty** | **DMSO** | **GSK-J4 Hydrochloride** | **UNC0638** | **A366** | **GSK J1** | **MS436** | **A196** | **Tasquinimod** | **IU1** | **AR42** | **Empty** |
| **Empty** | **AR42** | **UNC0642** | **UNC1999** | **EPZ6438** | **MI-3** | **PAOA** | **OICR-9429** | **TMP 195** | **Cambinol** | **DMSO** | **Empty** |
| **Empty** | **AR42** | **UNC1215** | **IOX1** | **EPZ004777** | **(R)-PFI-2 hydrochloride** | **RG2833** | **EPZ011989** | **EED226** | **GSK2879552 2HCl (1401966-69-5(free base))** | **DMSO** | **Empty** |
| **Empty** | **AR42** | **ML324** | **JIB04** | **CPI203** | **SGC707** | **Gambogenic acid** | **I-BRD9** | **LMK235** | **CPI0610** | **DMSO** | **Empty** |
| **Empty** | **Empty** | **Empty** | **Empty** | **Empty** | **Empty** | **Empty** | **Empty** | **Empty** | **Empty** | **Empty** | **Empty** |

| **Empty** | **Empty** | **Empty** | **Empty** | **Empty** | **Empty** | **Empty** | **Empty** | **Empty** | **Empty** | **Empty** | **Empty** |
| --- | --- | --- | --- | --- | --- | --- | --- | --- | --- | --- | --- |
| **Empty** | **DMSO** | **CB5083** | **Pyroxamide** | **Pimelic diphenylamide 106** | **Bromosporine** | **ORY1001** | **Delanzomib** | **(+)-JQ-1** | **MS049 2HCl (1502816-23-0(free base))** | **AR42** | **Empty** |
| **Empty** | **DMSO** | **ML240** | **5-Azacytidine** | **Ixazomib** | **Birabresib** | **Selisistat** | **TCS PIM-1 1** | **Histone Deacetylase Inhibitor III** | **P22077** | **AR42** | **Empty** |
| **Empty** | **DMSO** | **SRT2104** | **Vorinostat** | **BRD3308** | **RGFP 966** | **Tubastatin A** | **Procainamide hydrochloride** | **BIX-01294** | **LDN57444** | **AR42** | **Empty** |
| **Empty** | **AR42** | **Bobcat339 hydrochloride** | **Bortezomib** | **SantacruzaMate A** | **GSK2801** | **PCI34051** | **Zebularine** | **BIX-01294** | **ML323** | **DMSO** | **Empty** |
| **Empty** | **AR42** | **Mivebresib** | **Carfilzomib** | **ACY738** | **GSK591** | **VR23** | **Givinostat hydrochloride monohydrate** | **SIS17** | **Empty** | **DMSO** | **Empty** |
| **Empty** | **AR42** | **ORY-1001(trans)** | **Molibresib** | **ACY-775** | **CPI455** | **SGI1027** | **Chidamide** | **NMS 873** | **Empty** | **DMSO** | **Empty** |
| **Empty** | **Empty** | **Empty** | **Empty** | **Empty** | **Empty** | **Empty** | **Empty** | **Empty** | **Empty** | **Empty** | **Empty** |

**Supplementary Table 3: Plate view of the function of ordered compounds from MSD compound library (3 plates used for compound screening).**

| **Empty** | **Empty** | **Empty** | **Empty** | **Empty** | **Empty** | **Empty** | **Empty** | **Empty** | **Empty** | **Empty** | **Empty** |
| --- | --- | --- | --- | --- | --- | --- | --- | --- | --- | --- | --- |
| **Empty** | **DMSO** | **HDMi** | **HDACi** | **Epigenetic Reader Domain inhibitor; Others** | **GABA Receptor activator; Sodium Channel inhibitor; HDACi** | **Adrenergic Receptor; Dopamine Receptor; HMTi** | **Epigenetic Reader Domain** | **Sirtuin** | **Sirtuin inhibitor** | **AR42** | **Empty** |
| **Empty** | **DMSO** | **NF-κB inhibitor; HDAC** | **HDMi** | **HDACi** | **HDMi** | **Epigenetic Reader Domain inhibitor** | **HDACi** | **Epigenetic Reader Domain inhibitor** | **DUB** | **AR42** | **Empty** |
| **Empty** | **DMSO** | **Sirtuin inhibitor** | **HDACi** | **HDACi** | **Sirtuin activator** | **Proteasome inhibitor** | **HDAC** | **HDACi** | **HMTi** | **AR42** | **Empty** |
| **Empty** | **AR42** | **HDACi** | **HDACi** | **RAAS; HDACi** | **HMTi** | **DNMTi** | **HDACi** | **p97 inhibitor** | **HMTi** | **DMSO** | **Empty** |
| **Empty** | **AR42** | **Proteasome agonist** | **Sirtuin inhibitor** | **AChR antagonist; HDAC activator; PDE inhibitor** | **DUB inhibitor** | **DNMTi; DUB** | **HDACi** | **Epigenetic Reader Domain inhibitor** | **DNMTi** | **DMSO** | **Empty** |
| **Empty** | **AR42** | **HDAC activator** | **HDACi** | **Endogenous Metabolite; DNMTi** | **Adenosine Receptor antagonist; HDACi; PDE inhibitor; PKA activator; TNF inhibitor** | **HDACi** | **HDACi** | **DNMTi; HMTi** | **Epigenetic Reader Domain inhibitor** | **DMSO** | **Empty** |
| **Empty** | **Empty** | **Empty** | **Empty** | **Empty** | **Empty** | **Empty** | **Empty** | **Empty** | **Empty** | **Empty** | **Empty** |

| **Empty** | **Empty** | **Empty** | **Empty** | **Empty** | **Empty** | **Empty** | **Empty** | **Empty** | **Empty** | **Empty** | **Empty** |
| --- | --- | --- | --- | --- | --- | --- | --- | --- | --- | --- | --- |
| **Empty** | **DMSO** | **HDACi** | **Epigenetic Reader Domain inhibitor** | **HDMi** | **Epigenetic Reader Domain inhibitor** | **HMTi** | **Epigenetic Reader Domain inhibitor** | **HMTi** | **Proteasome inhibitor** | **AR42** | **Empty** |
| **Empty** | **DMSO** | **Epigenetic Reader Domain inhibitor** | **HMTi** | **HDMi** | **HMTi** | **Epigenetic Reader Domain inhibitor** | **Epigenetic Reader Domain** | **HMTi** | **DUB inhibitor** | **AR42** | **Empty** |
| **Empty** | **DMSO** | **HDMi** | **HMTi** | **HMTi** | **HDMi** | **Epigenetic Reader Domain inhibitor** | **HMTi** | **HDACi** | **DUB inhibitor** | **AR42** | **Empty** |
| **Empty** | **AR42** | **HMTi** | **HMTi** | **HMTi** | **Apoptosis; Epigenetic Reader Domain; HMTi** | **HDACi** | **HMTi ; JAK** | **HDACi** | **Sirtuin inhibitor** | **DMSO** | **Empty** |
| **Empty** | **AR42** | **Epigenetic Reader Domain antagonist** | **HDMi** | **HMTi** | **HMTi** | **HDACi** | **HMTi** | **DNMTi ; HMTi** | **HDMi** | **DMSO** | **Empty** |
| **Empty** | **AR42** | **HDMi** | **HDMi; HMTi** | **Epigenetic Reader Domain inhibitor** | **HMTi** | **HMTi** | **Epigenetic Reader Domain** | **HDACi** | **c-Myc inhibitor; Epigenetic Reader Domain inhibitor** | **DMSO** | **Empty** |
| **Empty** | **Empty** | **Empty** | **Empty** | **Empty** | **Empty** | **Empty** | **Empty** | **Empty** | **Empty** | **Empty** | **Empty** |

| **Empty** | **Empty** | **Empty** | **Empty** | **Empty** | **Empty** | **Empty** | **Empty** | **Empty** | **Empty** | **Empty** | **Empty** |
| --- | --- | --- | --- | --- | --- | --- | --- | --- | --- | --- | --- |
| **Empty** | **DMSO** | **p97 inhibitor** | **HDACi** | **HDACi** | **Epigenetic Reader Domain;CDK** | **HDMi** | **proteasome inhibitor** | **Epigenetic Reader Domain** | **HMTi** | **AR42** | **Empty** |
| **Empty** | **DMSO** | **p97 inhibitor** | **DNMTi** | **Proteasome inhibitor; Caspase** | **Epigenetic Reader Domain inhibitor** | **Sirtuin** | **Pim inhibitor** | **HDACi** | **DUB inhibitor** | **AR42** | **Empty** |
| **Empty** | **DMSO** | **Sirtuin activator** | **HDACi** | **HDACi** | **HDACi** | **HDACi** | **AChR inhibitor; DNMTi inhibitor;** | **HMTi** | **DUB inhibitor** | **AR42** | **Empty** |
| **Empty** | **AR42** | **DNMTi** | **Proteasome** | **HDACi** | **Epigenetic Reader Domain inhibitor** | **HDACi** | **DNMTi** | **HMTi** | **DUB inhibitor** | **DMSO** | **Empty** |
| **Empty** | **AR42** | **Epigenetic Reader Domain inhibitor** | **Proteasome inhibitor** | **HDACi** | **HMTi** | **Caspase inhibitor; proteasome inhibitor** | **HDACi** | **HDACi** | **Empty** | **DMSO** | **Empty** |
| **Empty** | **AR42** | **HDMi** | **Reader inhibitor** | **HDACi** | **HDMi** | **DNMTi** | **HDACi** | **p97 inhibitor** | **Empty** | **DMSO** | **Empty** |
| **Empty** | **Empty** | **Empty** | **Empty** | **Empty** | **Empty** | **Empty** | **Empty** | **Empty** | **Empty** | **Empty** | **Empty** |

**Supplementary Table 4: Plate view of targets of ordered compounds from MSD compound library (3 plates used for compound screening).**

| **Empty** | **Empty** | **Empty** | **Empty** | **Empty** | **Empty** | **Empty** | **Empty** | **Empty** | **Empty** | **Empty** | **Empty** |
| --- | --- | --- | --- | --- | --- | --- | --- | --- | --- | --- | --- |
| **Empty** | **DMSO** | **HDM** | **HDAC1; HDAC2; HDAC3** | **p300 HAT; KEAP1-Nrf2** | **GABA; Sodium Channel; HDACs; HDAC1** | **Adrenergic Receptor; D2; D3; Dopamine; α2-adrenergic** | **PCAF (p300/CBP-associated factor); p300 HAT; p300/CBP** | **SIRT1** | **SIRT1** | **AR42** | **Empty** |
| **Empty** | **DMSO** | **NF-κB** | **PRMT1; yeast Hmt1p** | **HDAC** | **KDM2/7 JmjC** | **p300** | **HDAC** | **BD2** | **USP7** | **AR42** | **Empty** |
| **Empty** | **DMSO** | **SIRT1; SIRT2** | **HDAC** | **HDAC** | **SIRT** | **Dipeptidase 1** | **HDAC** | **HDAC10; HDAC3; HDAC6; HDAC8; HDAC9** | **PRMT1** | **AR42** | **Empty** |
| **Empty** | **AR42** | **HDAC** | **HDAC** | **ACE; HDAC** | **HDMs** | **DNMTs** | **HDAC** | **p97** | **SMYD3** | **DMSO** | **Empty** |
| **Empty** | **AR42** | **Protease-Activated Receptor 2** | **SIRT** | **Adenosine receptor; HDAC2; PDE** | **JOSD2; SENP6 core; UCH-L3; USP4; USP8** | **DNMT1; USP2** | **HDAC** | **BRPF1B; BRPF2** | **DNMT** | **DMSO** | **Empty** |
| **Empty** | **AR42** | **HDAC** | **HDAC1; HDAC3** | **Human Endogenous Metabolite; DNMT** | **A1; A2; A3; HDAC2; PDE4; PKA; TNF-α** | **HDAC8; HDAC1;HDAC6** | **HDAC1; HDAC11; HDAC2; HDAC3** | **DNMT1; PRMT1** | **p300/CBP** | **DMSO** | **Empty** |
| **Empty** | **Empty** | **Empty** | **Empty** | **Empty** | **Empty** | **Empty** | **Empty** | **Empty** | **Empty** | **Empty** | **Empty** |

| **Empty** | **Empty** | **Empty** | **Empty** | **Empty** | **Empty** | **Empty** | **Empty** | **Empty** | **Empty** | **Empty** | **Empty** |
| --- | --- | --- | --- | --- | --- | --- | --- | --- | --- | --- | --- |
| **Empty** | **DMSO** | **HDAC1; HDAC2; HDAC3; HDAC6; HDAC8** | **CREBBP; EP300** | **LSD1** | **BRD2; BRD3; BRD4** | **PRMT5** | **L3MBTL1** | **PRMT1; PRMT3; PRMT4; PRMT6; PRMT8** | **Dipeptidyl peptidase II** | **AR42** | **Empty** |
| **Empty** | **DMSO** | **BRD2; BRD4** | **EZH1; EZH2** | **LSD1** | **SMYD2** | **BRPF1; BRPF2; BRPF3** | **BRD9** | **Coactivator Associated Arginine Methyltransferase 1 (CARM1)** | **UCH-L5** | **AR42** | **Empty** |
| **Empty** | **DMSO** | **JMJD3** | **G9a/GLP** | **G9a/GLP** | **JMJD3 (KDM6B);UTX (KDM6A)** | **BRD4 (1); BRD4 (2)** | **SUV420H1;SUV420H2** | **HDAC4** | **USP14** | **AR42** | **Empty** |
| **Empty** | **AR42** | **G9a/GLP** | **EZH1; EZH2** | **EZH2; EZH2** | **Apoptosis; Epigenetic Reader Domain; Menin-MLL** | **HDAC** | **HMT; WDR5** | **HDAC4; HDAC5; HDAC7; HDAC9** | **SIRT1; SIRT2** | **DMSO** | **Empty** |
| **Empty** | **AR42** | **L3MBTL3; L3MBTL3; L3MBTL3-D274A** | **2OG oxygenases, JmjC demethylases; KDM4A/3A** | **DOT1L** | **SETD7** | **HDAC1; HDAC3** | **EZH2** | **EED; PRC2** | **LSD1** | **DMSO** | **Empty** |
| **Empty** | **AR42** | **JMJD2** | **JMJD2A; JMJD2B; JMJD2D; JMJD2E; JARID1A** | **BRD4** | **PRMT3; PRMT3** | **EZH2** | **BRD9** | **HDAC4; HDAC5** | **MYC; BRD4-BD1** | **DMSO** | **Empty** |
| **Empty** | **Empty** | **Empty** | **Empty** | **Empty** | **Empty** | **Empty** | **Empty** | **Empty** | **Empty** | **Empty** | **Empty** |

| **Empty** | **Empty** | **Empty** | **Empty** | **Empty** | **Empty** | **Empty** | **Empty** | **Empty** | **Empty** | **Empty** | **Empty** |
| --- | --- | --- | --- | --- | --- | --- | --- | --- | --- | --- | --- |
| **Empty** | **DMSO** | **p97** | **HDAC** | **HDAC** | **BRD; CECR2** | **LSD1/KDM1A** | **Chymotrypsin-like proteasome** | **BRD4** | **PRMT4; PRMT6** | **AR42** | **Empty** |
| **Empty** | **DMSO** | **P97** | **DNMT** | **20S proteasome; 20S** | **BRDs** | **SIRT1** | **Pim1** | **HDAC** | **USP2; USP4; USP47; USP5; USP7** | **AR42** | **Empty** |
| **Empty** | **DMSO** | **SIRT1** | **HDAC1, HDAC2; HDAC3; HDAC6; HDAC8** | **HDAC3** | **HDAC3** | **HDAC6** | **AChR; DNMT1; Sodium Channel** | **G9a** | **UCH-L1; UCH-L1; UCH-L3** | **AR42** | **Empty** |
| **Empty** | **AR42** | **TET1; TET2** | **20S proteasome** | **HDAC2; HDAC4; HDAC6** | **BAZ2A; BAZ2B** | **HDAC1; HDAC10; HDAC2; HDAC6; HDAC8** | **Cytidine deaminase** | **G9a** | **USP1-UAF1** | **DMSO** | **Empty** |
| **Empty** | **AR42** | **BET** | **Proteasome** | **HDAC6** | **PRMT5** | **Caspase/trypsin/chemotrypsin like proteasomes** | **HD1-A; HD1-B; HD2** | **HDAC11** | **#N/A** | **DMSO** | **Empty** |
| **Empty** | **AR42** | **LSD1/KDM1A** | **BET proteins** | **HDAC6** | **KDM5A** | **DNMT1; DNMT3A; DNMT3B** | **HDAC1; HDAC2; HDAC3; HDAC8** | **p97** | **#N/A** | **DMSO** | **Empty** |
| **Empty** | **Empty** | **Empty** | **Empty** | **Empty** | **Empty** | **Empty** | **Empty** | **Empty** | **Empty** | **Empty** | **Empty** |

**Supplementary Table 5: List of additional purchased compounds for dose-response curve testing.**

| **Compounds** | **Supplier** | **Catalogue Number** | **Other information** |
| --- | --- | --- | --- |
| **WDR5-0103** | **MedChemExpress** | **HY-19347** |  |
| **GSK-LSD1** | **MedChemExpress** | **HY-100546** |  |
| **AR-42** | **MedChemExpress** | **HY-13265** |  |
| **MI-503** | **MedChemExpress** | **HY-16925** |  |
| **BAY598** | **MedChemExpress** | **HY-19546** |  |
| **PBIT** | **MedChemExpress** | **HY-101451** |  |
| **MI-463** | **MedChemExpress** | **HY-19809** |  |
| **MI-538** | **MedChemExpress** | **HY-19810** |  |
